# Effects of abnormal muscle forces on prenatal joint morphogenesis in mice

**DOI:** 10.1101/669267

**Authors:** Vivien Sotiriou, Rebecca A Rolfe, Paula Murphy, Niamh C Nowlan

**Affiliations:** Department of Bioengineering, Imperial College London, UK; Department of Zoology, School of Natural Sciences, Trinity College Dublin, The University of Dublin, Ireland

**Author notes:** Corresponding author: Dr Niamh C Nowlan, Department of Bioengineering, Royal School of Mines, Prince Consort Road, Imperial College London, SW7 2AZ, United Kingdom. Author Contribution: NCN and PM conceived the idea for this project, RR collected the embryonic murine limb scan data. PS analysed the data and interpreted the results. PS drafted the manuscript and NCN, PM and RR critically revised it. All authors gave final approval of the manuscript to be published.

**Keywords:** Fetal movements, biomechanics, image registration, mouse embryos, muscleless limbs, reduced muscle, joint development, joint shape

## Abstract

Fetal movements are essential for normal development of the human skeleton. When fetal movements are reduced or restricted, infants are at higher risk of developmental dysplasia of the hip and arthrogryposis (multiple joint contractures). Joint shape abnormalities have been reported in mouse models with abnormal or absent musculature, but the effects on joint shape in such models have not been quantified or characterised in detail. In this study, embryonic mouse forelimbs and hindlimbs at a single developmental stage (Theiler Stage 23) with normal, reduced or absent muscle were imaged in 3D. Skeletal rudiments were virtually segmented and rigid image registration was used to reliably align rudiments with each other, enabling repeatable assessment and measurement of joint shape differences between normal, reduced-muscle and absent muscle groups. We demonstrate qualitatively and quantitatively that joint shapes are differentially affected by a lack of, or reduction in, skeletal muscle, with the elbow joint being the most affected of the major limb joints. Surprisingly, the effects of reduced muscle were often more pronounced than those of absent skeletal muscle, indicating a complex relationship between muscle mass and joint morphogenesis. These findings have relevance for human developmental disorders of the skeleton in which abnormal fetal movements are implicated, particularly developmental dysplasia of the hip and arthrogryposis.

## Introduction

Fetal movements are important for the development of healthy joints reviewed in ^1, 2^, and absent or abnormal fetal movements can lead to developmental skeletal abnormalities such as developmental dysplasia of the hip (DDH) ^1, 3^ and arthrogryposis ^4^. DDH is a condition in which the immature hip joint is unstable or dislocated ^5^, with an incidence of 1.3 per 1000 live births ^6^. While a positive family history and female gender are risk factors associated with the condition, the other major risk factors relate to the freedom of movement of the fetus. For example, fetal breech position ^7^ and oligohydramnios (a reduction of amniotic fluid) ^8^ increase the risk of DDH. Arthrogryposis, a syndrome with multiple joint contractures, is another developmental disorder associated with abnormal muscle activity and thus abnormal movement *in utero* ^9^. While a range of different genetic abnormalities may lead to arthrogryposis ^4^, abnormal fetal movements are common to all manifestations of the syndrome reviewed in ^1^. Despite the clinical significance of fetal movements in joint morphogenesis, their contribution to the emergence of shape of different joint types has not yet been quantified.

While the very early stages of joint development occur independently of fetal movements ^10, 11^, numerous animal studies have described the importance of movement for morphogenesis and cavitation of the joint reviewed in ^2,12,13,14^. The most commonly used model systems are pharmacologically immobilised chick and zebrafish embryos, zebrafish mutants and genetically modified mouse lines in which skeletal muscle is reduced, absent or abnormal ^15−19^. Immobilised joints of developing chicks and mice fail to undergo cavitation ^2,3,10,13,15^, while a range of shape effects have been reported in zebrafish, chicks and mice including simpler, smaller or malformed articular surfaces with absent or reduced muscle attachment sites ^13, 15, 16, 18, 19.^ We previously showed that when fetal movements are absent in the chick, early joint morphogenesis proceeds as normal up until the point at which joint cavitation should occur, but that when cavitation does not occur, subsequent shape development of the hip joint is abnormal ^3^. There is an intriguing difference in skeletal development between pharmacologically immobilised chicks and so-called “muscleless limb” genetically modified mice. When spontaneous fetal movements are absent in chicks, all bones and synovial joints seem to be affected to a similar degree reviewed in ^12^, while in immobile mouse embryos, the rudiments of the forelimb are more severely affected than those of the hindlimb ^15^. For example, cavitation and shape of the murine elbow joint is substantially disrupted by the absence of skeletal muscle ^10, 15^, while the shape of the knee joint does not show obvious abnormalities ^15^. We have previously proposed that passive movements induced by activities of the mother and healthy littermates may be compensating to some degree for the lack of spontaneous movements in muscleless limb mice, with variable effects in different regions of the limbs ^20^.

Although attempts to quantify shape change due to immobility have been made for the chick knee joint ^13, 21^, no extensive quantitative assessment has been performed for multiple joints, or for any joints of the paralysed developing mouse. It is challenging to perform consistent, repeatable measurements on complex 3D shapes such as synovial joints, because it is very difficult to ensure precise alignment prior to taking measurements. Image registration is an approach that can resolve this challenge ^22^. In this study, we perform rigid image registration on 3D segmentations of long bone rudiments from genetically modified mouse embryos with normal, reduced (*Myf5*^*nlacZ/+:*^*Myod*^*−/−*^), or absent limb muscle (*Myf5*^*nlacZ/nlacZ*^*:Myod*^*−/−*^), to investigate the effects of muscle volume on the shape of the elbow, shoulder, hip and knee joints. We focus on Theiler Stage (TS ^23^) 23 at approximately embryonic day (E) 14.5, when statistically significant effects on ossification due to reduced or absent skeletal muscle are apparent in some rudiments but not in others ^15^. The aim of this study was to use image registration to reliably align rudiments of normal, reduced-muscle and muscleless limbs, to systematically quantify the effects of reduced or absent muscle on the shapes of the major synovial joints.

## Methods

### Animal model and 3D data acquisition

The data used for this study is a subset of the data described but not quantitatively analysed for joint shape in a previous study ^15^ where double heterozygous Myf5^nlacZ/+^:Myod^+/−^ mice were interbred to generate E14.5 or E15.5 embryos, staged as developmental stage TS23. The control genotypes included; wildtype, *Myf5*^*nlacZ/+:*^*Myod*^*+/−*^, *Myf5*^*nlacZ/+*^*:Myod*^*+/+*^ *and Myf5*^*+/+*^*:Myod*^*−/+*^. The reduced-muscle and the muscleless genotypes were *Myf5*^*nlacZ/+:*^*Myod*^*−/−*^ and *Myf5*^*nlacZ/nlacZ:*^*Myod*^*−/−*^ respectively. Limbs were dissected, stained with Alcian blue and Alizarin red, and scanned with Optical Projection Tomography as described previously ^15^. 3D data of reconstructed hindlimbs and forelimbs were used for analysis. All animal work followed the guidelines of the Trinity College Dublin Bioresources Unit and Bioethics Committee.

### Image registration

Each limb reconstruction was imported into Mimics (Materialise, Leuven, Belgium). A minimum of five forelimbs and hindlimbs for each of the three groups (control, reduced-muscle, muscleless) were successfully segmented and analysed, as reported in Table 1. The scapula, humerus, radius and ulna were individually segmented for each forelimb, and the pelvis, femur and tibia for the hindlimb. It was possible to reliably segment individual rudiments even when the joints appeared to be fused, due to consistently reduced brightness in the failed joint line. For each experimental group, segmented rudiments of the same type (e.g., all humeri) were orientated similarly using the ImageJ plugin TransformJ (http://rsbweb.nih.gov/ij/, last accessed October 2018) ^24, 25^. Image brightness within each group was normalised with a MATLAB algorithm (version R2015a, The MathWorks, Inc., Natick, Massachusetts, United States). One rudiment in each group was selected as the target image onto which others were orientated. All other rudiments were then rigidly registered to the target and an “atlas” was created for each rudiment (scapula, humerus, radius, ulna) and each group (control, reduced-muscle, muscleless) with the use of the Image Registration Toolkit software ^26^ (IRTK, BioMedia, Imperial College London), as described previously ^27^. Further rigid registrations were performed to precisely align each of the three rudiment atlases with each other, with the same manipulations then being applied to each individual rudiment dataset. At the end of this process, each individual rudiment was precisely aligned with all other rudiments for the three experimental groups. With seven rudiments interfacing at the joints of interest and three experimental groups, a total of 21 atlases were created. To visualise the differences in shape and size of the articular surfaces between groups, control, reduced-muscle and muscleless atlases for each rudiment were overlaid using ImageJ (atlas videos included in supplementary material: videos S1-S7).

**Table 1:** Numbers of limbs analysed in this study for the three groups.

|  | Control | Reduced-muscle | Muscleless |
| --- | --- | --- | --- |
| Forelimb | 5 | 5 | 5 |
| Hindlimb | 5 | 5 | 7 |

### Qualitative and quantitative analysis of data

Regions of pronounced shape changes between groups were identified by applying repeatable, recorded rotations to the atlas overlays, and measurements defined to quantitatively assess these apparent differences as shown in Figures 1 and 2. Consistent measurements could be made from equivalent sections and planes of each individual rudiment by applying the same rotations used for the atlases. The planar orientations used are shown in Figure 3. Measurements were performed in Gwyddion image editing software (Gwyddion 2.44, http://gwyddion.net/, last accessed October 2018). As differences in rudiment length were previously reported for some reduced-muscle and muscleless rudiments ^15^, measurements were normalised by the length of the rudiment under investigation, in order to focus the outcomes on shape-specific (rather than overall size dependent) changes. One-way analysis of variance tests (ANOVAs) (SPSS Statistics 24, IBM corp., Armonk, NY), with significance level of α=0.05 were performed for all length-normalised measurements. Tukey’s post hoc test was used to identify pairwise differences between the three groups. If the variances between the groups were unequal, the Welch test was used and the pairwise differences were identified by the Games-Howell post hoc test. If the assumption for normality was not satisfied, the Kruskal-Wallis non-parametric test was used instead of ANOVAs. Only those results for which a significant difference (p-value<0.05) between at least two of the three groups was found are presented in detail and displayed graphically, while the full table of results is available in supplementary Table S8. Power analyses were performed for all 13 measurements with statistically significant differences and all had sufficient statistical power (>64%, power calculations in supplementary material S9). To allow for visual assessment of changes in shape between groups, rudiment shape outlines were traced on frontal, lateral and axial sections through the prime regions of interest for each rudiment.

**Figure 1:**
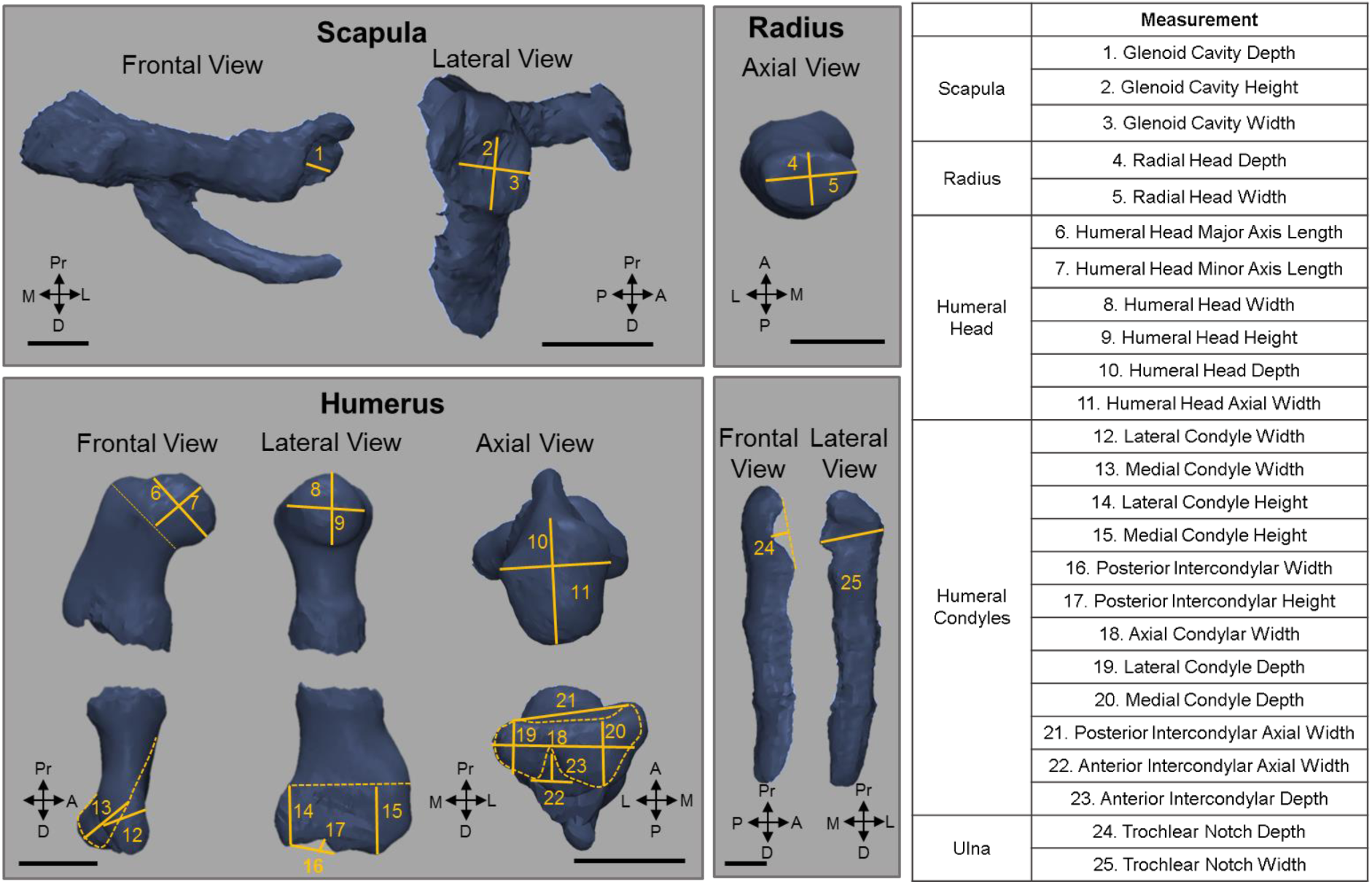
Measurements performed on individual rudiments of the forelimb. Left: locations of measurements (solid lines), with dashed lines indicating guides for measurements. Scale bars: 0.5mm. Pr: proximal, D: distal, A: anterior, P: posterior, M: medial, L: lateral. (Right) Reference numbers and nomenclature for all measurements performed.

**Figure 2:**
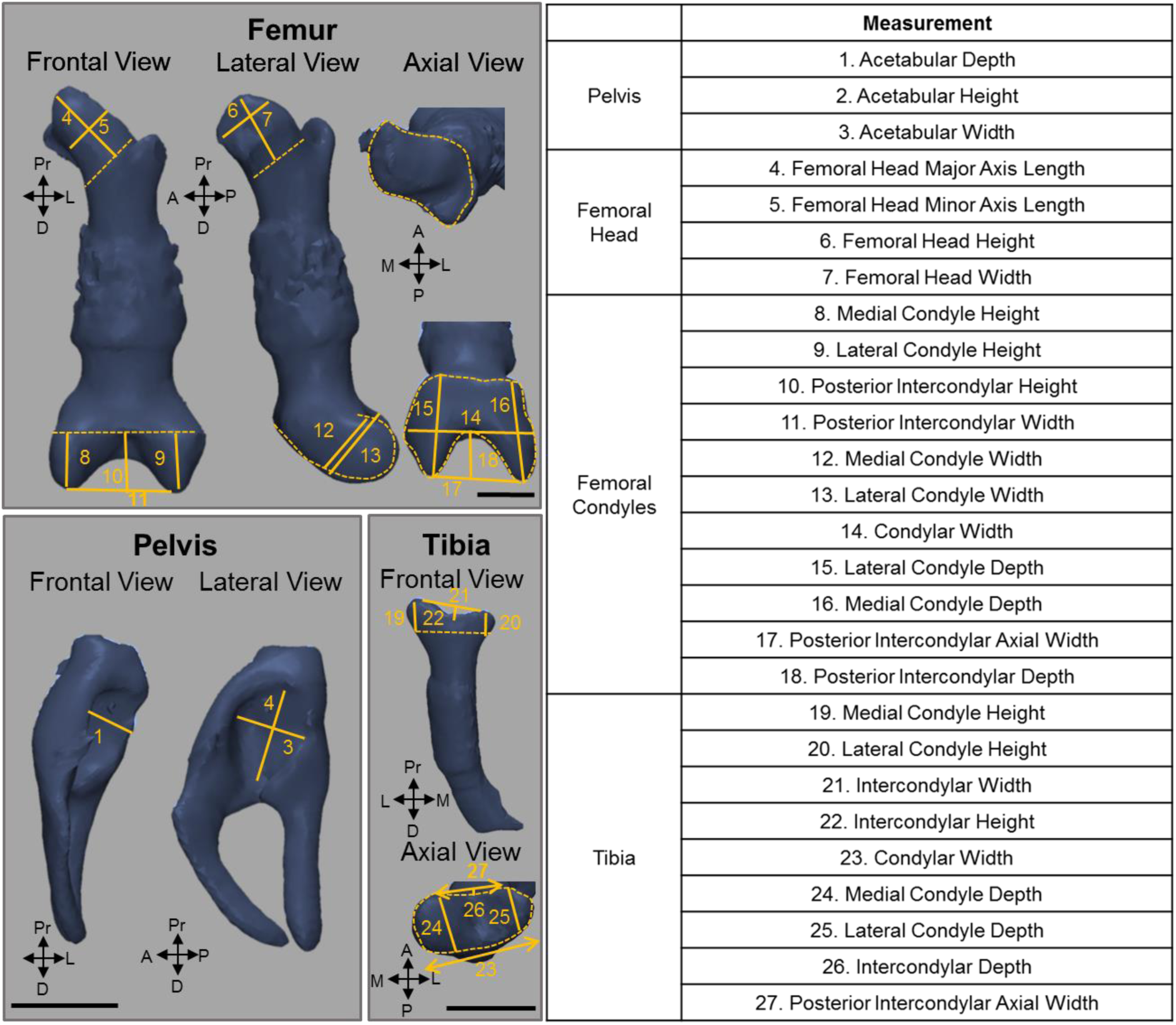
Measurements performed on individual rudiments of the hindlimb. Left: locations of measurements (solid lines), with dashed lines indicating guides for measurements. Scale bars: 0.5mm. Pr: proximal, D: distal, A: anterior, P: posterior, M: medial, L: lateral. Right: reference numbers and nomenclature for all measurements performed.

**Figure 3:**
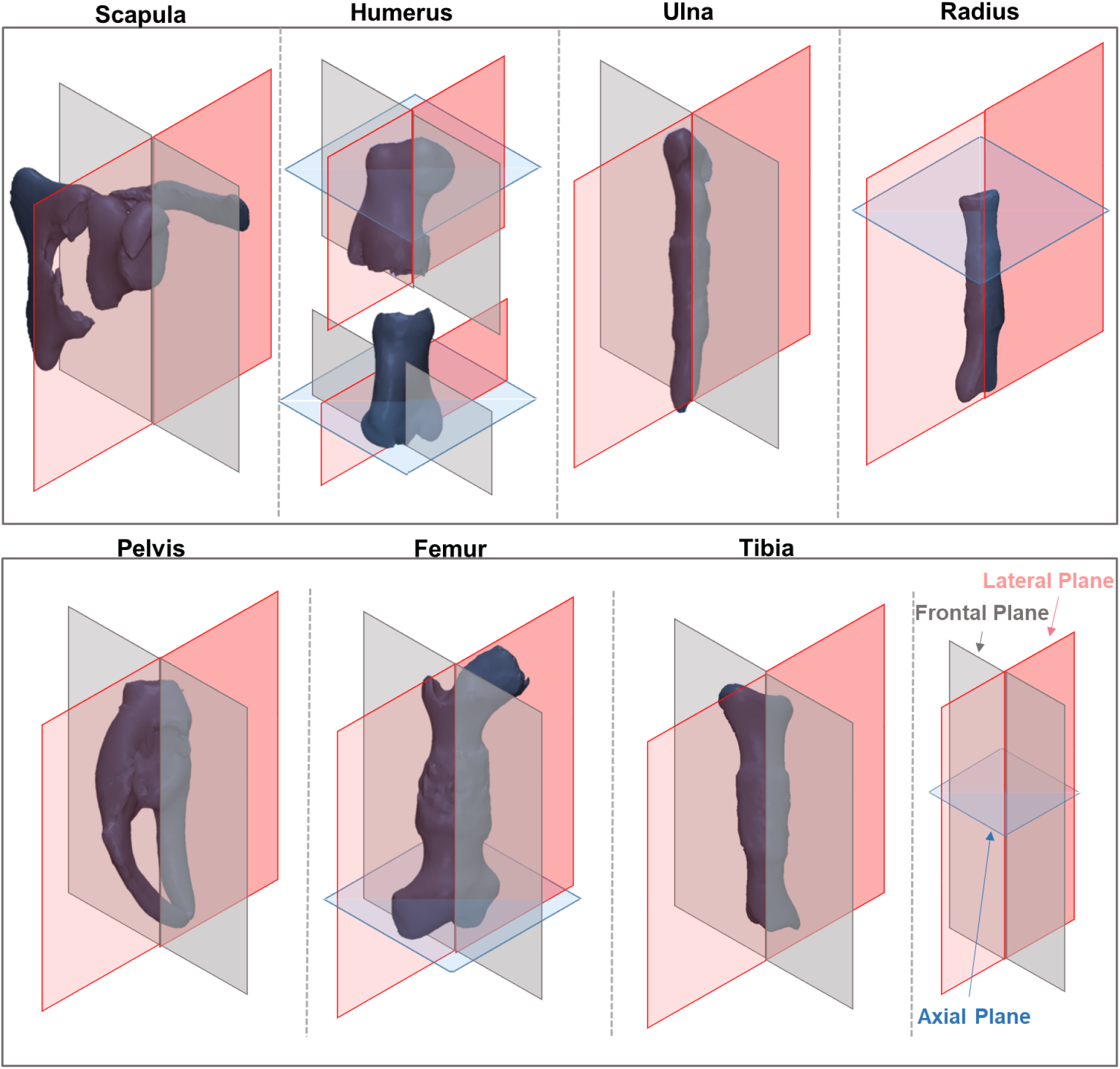
Planar orientations used for measurements of individual rudiments (not to scale).

## Results

The control, reduced-muscle and muscleless atlas overlays for each rudiment are provided in supplementary video files S1–S7, in which the control average shape is shown in blue, the reduced-muscle in yellow and the muscleless in purple. Bearing in mind that the atlases are based on the original (unscaled) data, the most obvious shape features are summarised below. There was a lack of separation evident between the coracoid process and the scapular body in the muscleless group (video S1), while a deformed shape was evident at the distal end of the muscleless humerus (video S2). Changes in the ulnar trochlear notch were evident in both the muscleless and reduced-muscle groups as compared to the controls (video S3). No apparent differences between the three groups were observed in the acetabular atlases (video S5), while the muscleless femoral head exhibited an abnormal protrusion in the region of the greater trochanter (video S6). In the proximal tibia, the depth of the intercondylar region varied between groups, with the reduced-muscle group appearing to have the shallowest intercondylar region and the controls having the deepest intercondylar region (video S7). The reduced-muscle tibiae appeared to have more prominent condyles than the control and muscleless tibiae (video S7). A detailed qualitative and quantitative assessment of shape for five major joints, informed by the atlas overlays, follows.

### Shoulder Joint

The most substantial shape changes in the reduced-muscle and muscleless shoulder joints were differences in the width and orientation of the glenoid cavities, which in the frontal plane appeared smaller in the reduced-muscle and muscleless limbs compared to the controls (Figure 4i:B, P-A orientation), and mal-aligned in the muscleless limbs (Figure 4i:A, dashed lines). However, the only significantly different measurement for the glenoid cavity was a reduction in depth (measurement 1, Figure 1) in the muscleless group compared to the controls (Figure 4ii). Differences in humeral head shape were also evident. They appeared to be flatter and wider in the reduced-muscle and muscleless groups (Figure 4i:C, D correspondingly), with the muscleless humeral heads missing the greater tubercle (Figure 4i:C, arrow). Quantitative differences in humeral head shape were found only for the reduced-muscle group, with humeral head height (measurement 9, Figure 1) significantly greater in the reduced muscle group than both other groups, and humeral head axial width (measurement 11, Figure 1) increased in the reduced-muscle group compared to the muscleless group (Figure 4ii).

**Figure 4:**
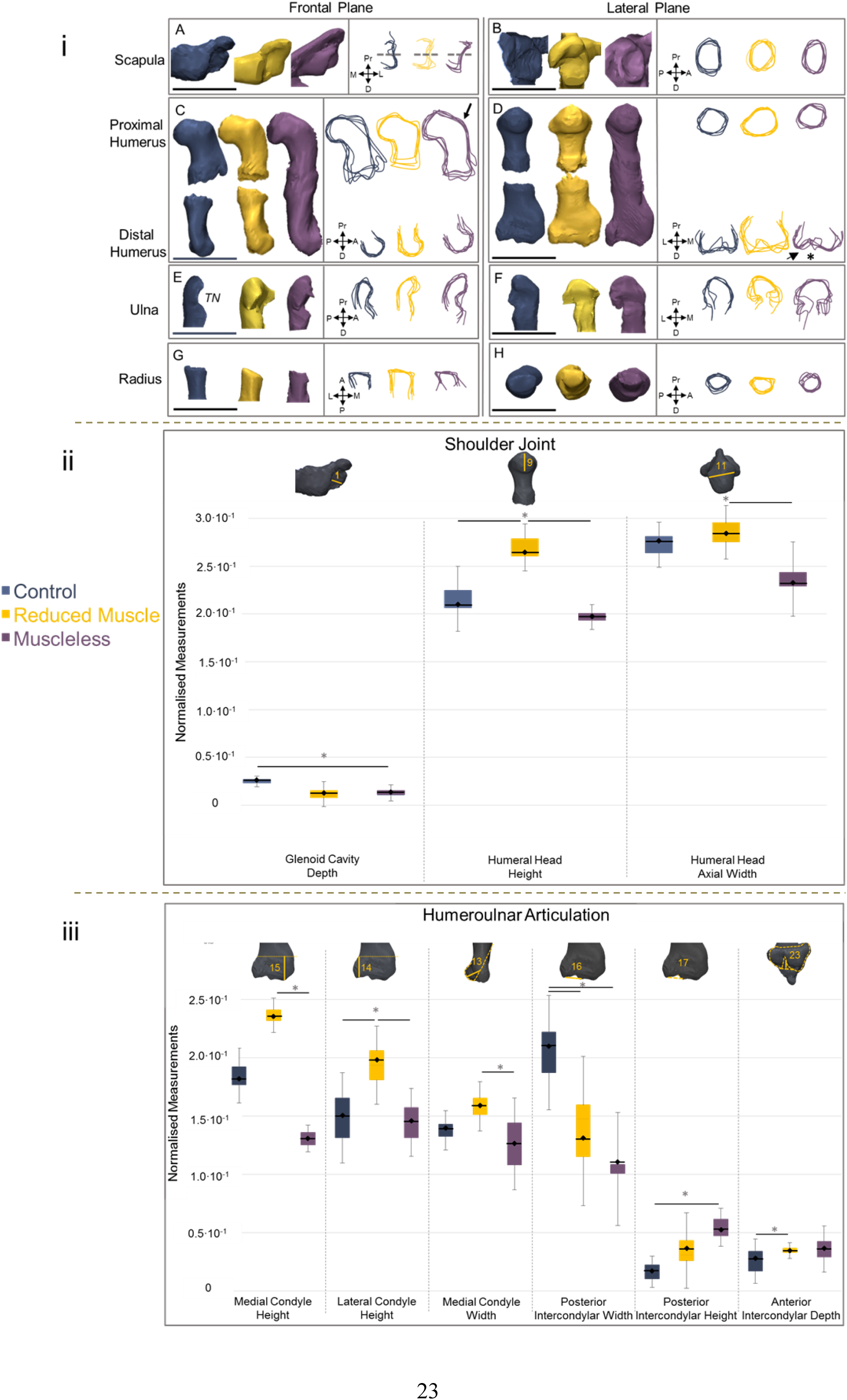
Abnormal muscle affects many aspects of elbow joint shape, with more limited effects on the shoulder joint. (i) Representative 3D shapes (left) and outlines of all five specimens (right) of control (blue), reduced-muscle (yellow) and muscleless (purple) forelimb rudiments. 3D shapes and outlines are of original sizes (i.e., not normalised). Key features of shape changes in the joints include: abnormal angle and increased height of the glenoid cavity in the muscleless group (A, B, purple outlines), abnormal prominence of the lateral condyle in the distal muscleless humerus (D, arrowhead) and greater variation in the shape of the proximal ulna in the reduced and muscleless groups (F, yellow and purple outlines). Pr: proximal, D: distal, A: anterior, P: posterior, M: medial, L: lateral, TN: Trochlear Notch. Scale bars: 1mm. (ii, iii) Length-normalised measurements (detailed in Figure 1) of the (ii) shoulder joint and (iii) humeroulnar joint for which significant (p value <0.05) differences between at least two groups were found.

### Elbow Joint

The key shape features evident from the 3D shapes and outlines of the distal humerus were wider condyles in the reduced-muscle limbs (Figure 4i:D), and an altered intercondylar region in the muscleless limbs (Figure 4i:D, asterisk). The muscleless specimens also exhibited an abnormally sharp protuberance of the lateral condyle (Figure 4i:D, arrow). Three condylar measurements and three measurements of the intercondylar region had significant differences between groups. For the posterior intercondylar width and height (measurements 16 & 17, Figure 1) the trends indicated progressive effects according to degree of muscle abnormality, with decreasing width and increasing height from control to reduced-muscle and from reduced-muscle to muscleless (Figure 4iii). Reduced-muscle and muscleless intercondylar widths were both significantly decreased compared to controls, while only the muscleless intercondylar height was significantly greater than controls (Figure 4iii). While anterior intercondylar depth also exhibited an increasing trend with muscle abnormality, only the reduced-muscle group was significantly increased compared to controls (Figure 4iii). In contrast, the condylar measurements (medial condyle width, and lateral and medial condyle height; measurements 13–15, Figure 1) of the reduced-muscle samples tended to be increased compared to both control and muscleless groups, with significant differences compared to one or both groups, and no significant differences between control and muscleless groups (Figure 4iii). Therefore, for the condyles of the distal humerus, the shape changes were divergent from what one might expect based on the assumption of a linear dependency on muscle volume. The reduced-muscle and muscleless proximal ulnae exhibited highly irregular shapes, with abnormal angulations visible in the lateral plane (Figure 4i:F). These differences were more pronounced in the muscleless group (Figure 4i:F, purple outlines). The shape and angle of the trochlear notch (Figure 4i:E, TN) also appeared affected in the reduced-muscle and muscleless groups with the convexity of the trochlear notch increasing from control to reduced-muscle to muscleless. However, there were no statistically significant differences between groups for ulnar measurements. No dramatic changes in shape of the proximal radius were evident (Figure 4i:G, H) and quantitative analyses of the humeroradial articulation showed no significant differences between control, reduced-muscle and muscleless groups. Therefore, in the elbow, the distal humerus exhibited more severe changes than its opposing rudiments.

### Hip Joint

Prominent qualitative differences in shape were observed in the muscleless acetabula and femoral heads (Figure 5i, A–D). A progressive loss of the normal shape of the acetabulum from control to reduced-muscle to muscleless pelvises was observed in the frontal plane (Figure 5i:A) with the shape of the muscleless acetabula resembling more a right angle than a concavity. Quantitative analysis revealed a trend of reduced acetabular width (measurement 3, Figure 2) from control to reduced-muscle to muscleless, with the only significant difference being between the muscleless group and the controls (Figure 5ii). The femoral heads of the reduced-muscle group appeared to be wider than those of the control or muscleless groups (Figure 5i:C, D), while the femoral heads of the muscleless group were noticeably irregular in the frontal plane, with a cylindrical profile in the lateral plane (Figure 5i:D, black arrowheads). The muscleless proximal femora also had an abnormally shaped outgrowth below the emerging greater trochanter (Figure 5i:C, asterisk). However, there were no significant differences between any of the three groups for measurements of the proximal femur. Therefore, in the muscleless group, the acetabulum was the most affected part of the hip joint with both qualitative and quantitative changes relative to the controls, with some shape changes visible qualitatively in the opposing femoral head.

**Figure 5:**
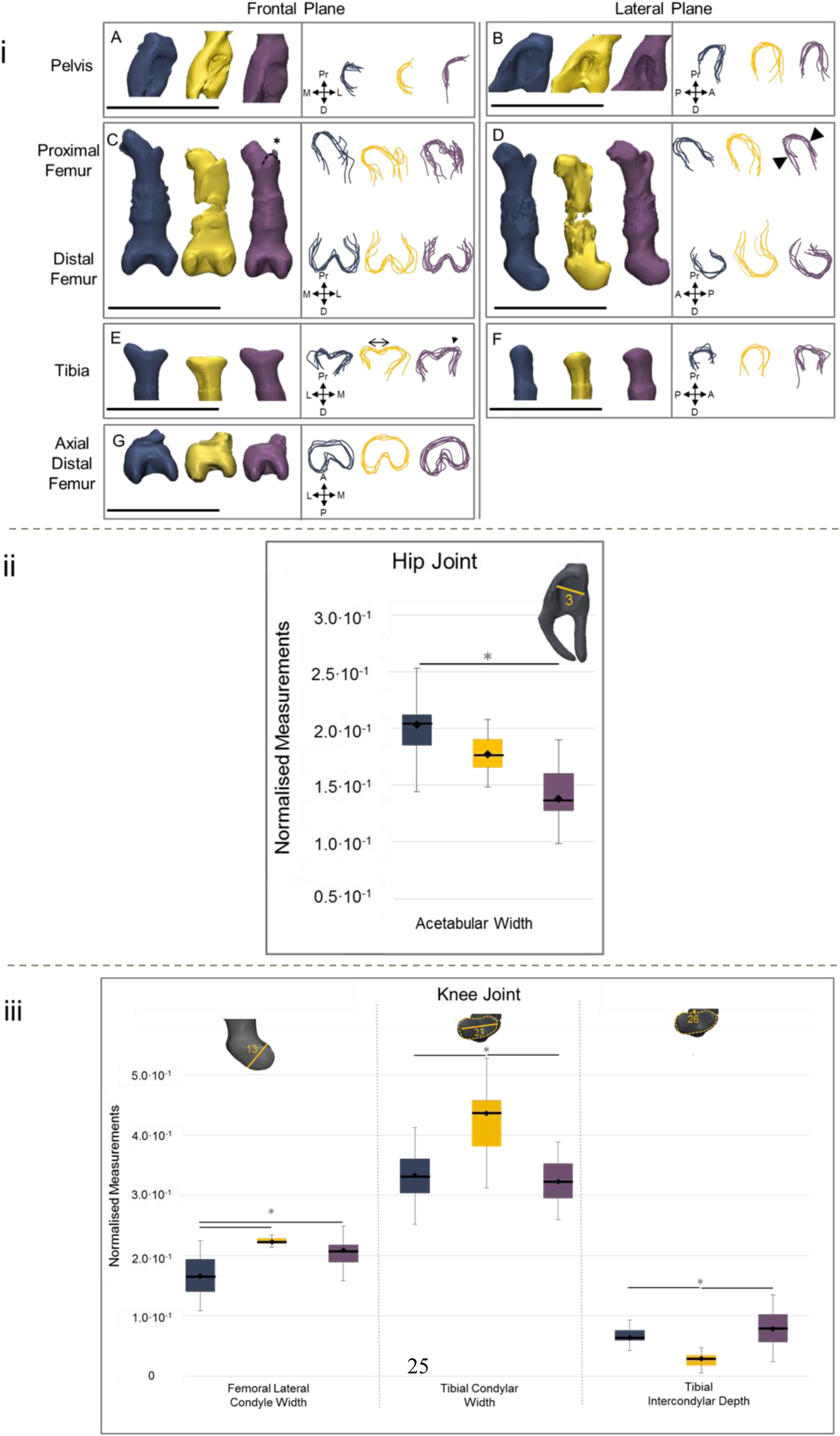
The knee and hip joint are subtly affected by abnormal muscle. (i) Representative 3D shapes and outlines of control (blue), reduced-muscle (yellow) and muscleless (purple) hindlimb rudiments. 3D shapes and outlines are of original sizes (i.e., not normalised). Key features of shape changes in the joints include: progressive loss of concavity of the acetabulum (A outlines), and increase in acetabular width, from control to muscleless rudiments (B, outlines), irregularity of the muscleless proximal femur (C, D), and abnormalities of the proximal tibia, increased width in the reduced-muscle group (E, double-headed arrow) and protrusion of the medial condyle of the muscleless proximal tibia (E, arrowhead). (ii, iii) Length-normalised measurements (detailed in Figure 2) of the (ii) hip joint and (iii) knee joint for which significant (p value <0.05) differences between at least two groups were found.

### Knee joint

There were few obvious shape abnormalities in the knee joints of the muscleless groups (Figure 5i:C–F), consistent with previous findings of a milder phenotype at the knee ^15^. However, the curvature of the femoral medial condyle appeared altered in the muscleless and reduced-muscle groups compared to the control group (Figure 5i:D), and quantitative analyses revealed significant increases in lateral condyle width (measurement 13, Figure 2) in the muscleless and reduced-muscle groups compared to controls (Figure 5iii). The reduced-muscle group appeared to have a wider tibial plateau than the other two groups (Figure 5i:E, double headed arrow), while the muscleless group had an abnormally sharp protuberance of the medial condyle (Figure 5i:E, arrowhead). The only significant differences found for the tibia were in the reduced-muscle group, with a significantly wider condylar region (measurement 23, Figure 2) and a significantly shallower intercondylar region (measurement 26, Figure 2) than both other groups (Figure 5iii). The shape of the knee joint was therefore slightly more severely affected in the reduced-muscle group than in the muscleless group.

## Discussion

In this study, the effects of absent and reduced skeletal muscle on emerging murine limb joint shape at one developmental stage were investigated. Image registration was used to precisely align the major skeletal rudiments of forelimbs and hindlimbs. The shapes of the shoulder, elbow, hip and knee joints were assessed qualitatively, using visual comparisons of aligned 3D renderings and 2D outlines, and quantitatively, using selected measurements of the component specimens. In line with previously published work ^10, 15^, we find that the elbow, hip and shoulder joints are affected by a complete absence of muscle, and report for the first time the effects on the shape of the knee joint. We also describe, for the first time, the effects on shape of a reduction in muscle volume. The elbow joint is the most severely affected by either absent or reduced muscle, with pronounced differences in shape and several significantly different measurements. Meanwhile, the muscleless shoulder, hip and knee have only one significantly different shape feature each compared to controls. A novel finding from this work is that, in contrast to the ‘dose-effect’ found for the amount of muscle on the ossification process in some rudiments (namely scapula and humerus) ^15^, the effects on joint shape are not linearly related to the amount of muscle. Indeed, in some joints, the shape changes in reduced-muscle rudiments are more pronounced than in the muscleless group. Finally, we find variation between joint types in the severity of effect between opposing surfaces of the same joint due to reduced or absent muscle. For example, in the elbow joint, the shape abnormalities of the distal humerus are not matched by similar effects in the opposing radius and ulna.

There are some limitations to this study. While our data was aligned in 3D, and regions of shape changes were identified using 3D data, qualitative and quantitative shape analysis was performed on 2D data sections. While we considered 3D shape quantification methods, such as principal component analysis (PCA) ^28^, our methods were preferable for visualising specific measurable differences between three distinct groups. With PCA, shapes from all groups must be grouped in one average shape. By keeping each individual original dataset (precisely aligned with all other rudiments of the same type), we were able to perform statistical analyses on specific, identifiable aspects of shape and anatomy. We were also able to highlight qualitative shape changes between the three groups which were not easily quantified. For example, we describe an increasingly irregular profile of the proximal ulna from control to reduced-muscle to muscleless groups, which was not evident from the quantitative measurements alone. The image registration and shape quantification techniques that we have described here are time consuming and not yet fully automated, and therefore may not be scalable to large datasets without further development of the methodological pipeline. Sample sizes are admittedly low, due to the difficulty of obtaining large numbers of mutant animals (double null embryos are produced at a rate of 1/16). However, power calculations demonstrated that all 13 significant differences had sufficient statistical power (as detailed in full in supplementary material, file S8) and therefore, we are confident that any results reported as being significantly different are indeed representative.

This study confirms previous observations on the loss of reciprocal and interlocking joint surfaces when muscle is absent, in the elbow, hip andshoulder ^10, 15^, but provides the first detailed and quantitative description of shape changes. While our previous work, based on observations of 3D reconstructions, stated that the knee joint “appeared unaffected” in the muscleless limb mice ^15^, this substantially more detailed analysis has revealed several shape changes in the knee joint due to the lack of muscle. This study provides the first description of the effects of reduced skeletal muscle on joint shape, and reveals a complex relationship between muscle volume and joint morphogenesis. The variation between joints for the effects of reduced muscle volume could be due to inherent differences between the joints or their dependence on muscle. It could also be that not all muscles are reduced by the same quantity in these mice, as muscle fibre volume for these mice has been quantified only in two muscles of the forelimb (the biceps and triceps branchii ^29^). Meanwhile, function or interaction of these abnormal muscles has not been quantified, which is another potentially confounding factor. For example, it is likely that the movements of the reduced-muscle mice are atypical, leading to abnormal (rather than simply reduced) mechanical stimulation of the joints. reviewed in ^302,10^

Our assessment of the relationship between interlocking joint surfaces offers an insight into how joint morphogenesis is directed, and the role of mechanical forces in this process. We find that when skeletal muscle is absent or reduced, opposing articulating surfaces are usually both abnormally shaped, but not always to the same degree. An influx model of joint development was recently proposed, in which cells continuously enter the interzone and help shape the joint ^31^. Shwartz et al ^31^ observed asymmetry in the contribution of Gdf-5 positive cells to opposing joint surfaces, and a previous paper from the same group demonstrated abnormal Gdf-5 expression in the interzone of a muscleless limb mouse model. Therefore, the asymmetry in shape effects between opposing articulating surfaces that we observe could be a downstream effect of abnormal modulation of mechanosensitive Gdf-5 progenitor cells. Another possible contributor to the asymmetry in shape effects observed is growth-related pressures and strains, as described by Henderson and Carter ^32^. As distinct from direct muscle forces or the biomechanical stimuli induced by movements, growth-related pressures and strains arise from internal forces and local deformations generated by differential growth rates. When skeletal muscle is present but with reduced volume, complex interactions between growth-related strains and pressures, muscle forces and fetal movement patterns could be leading to the varied outcomes seen between, and within, joints.

In conclusion, we precisely aligned the complex shapes of the developing mouse limb skeleton using image registration techniques, and used the aligned data to perform consistent measurements of the major synovial joints. We find that some joints are affected more than others, and that a reduction in limb muscle volume can have effects as severe as, or more pronounced than, the complete absence of muscle in some joints. These findings have relevance for human developmental disorders of the skeleton, particularly for developmental dysplasia of the hip and arthrogryposis, in which shape abnormalities are related in part to reduced, restricted, or abnormal fetal movements. It is still unclear why the hip joint is particularly dependant on fetal movements, or why there are particular joints which are more affected (in frequency or severity) than others with arthrogryposis ^9^. Our future work will apply the same techniques to different stages of development, to understand how joint shape abnormalities evolve over gestation.

## Acknowledgements

This research was funded by the European Research Council under the European Union’s Seventh Framework Programme (ERC Grant agreement no. [336306]). The generation of data analysed here was funded by a Wellcome Trust grant to PM (grant number 083539/Z/07/Z). The funders had no role in study design, data collection and analysis, decision to publish, or preparation of the manuscript. We are grateful to Shahragim Tajbakhsh and Gerard Dumas (Pasteur Institute, Paris, France) for providing embryonic material and Celine Bourdon for the processing of specimens previously reported^15^.

